# Activation of the mPFC-NAc pathway reduces motor impulsivity but does not affect risk-related decision-making in innately high-impulsive rats

**DOI:** 10.1101/2024.03.08.584121

**Authors:** Chloé Arrondeau, Ginna Urueña-Méndez, Florian Marchessaux, Raphaël Goutaudier, Nathalie Ginovart

## Abstract

Attention-deficit/hyperactivity disorder (ADHD) and substance use disorders (SUD) are characterized by exacerbated motor and risk-related impulsivities, which are associated with decreased cortical activity. In rodents, the medial prefrontal cortex (mPFC) and nucleus accumbens (NAc) have been separately implicated in impulsive behaviors, but studies on the specific role of the mPFC-NAc pathway in these behaviors are limited. Here, we investigated whether heightened impulsive behaviors are associated with reduced mPFC activity in rodents, and determined the involvement of the mPFC-NAc pathway in motor and risk-related impulsivities. We used the Roman High- (RHA) and Low-Avoidance (RLA) rat lines, which display divergent phenotypes in impulsivity. To investigate alterations in cortical activity in relation to impulsivity, regional brain glucose metabolism was measured using positron emission tomography and [^18^F]-fluorodeoxyglucose ([^18^F]FDG). Using chemogenetics, the activity of the mPFC-NAc pathway was either selectively activated in high-impulsive RHA rats or inhibited in low-impulsive RLA rats, and the effects of these manipulations on motor and risk-related impulsivity were concurrently assessed using the rat gambling task. We showed that basal [^18^F]FDG uptake was lower in the mPFC and NAc of RHA compared to RLA rats. Activation of the mPFC-NAc pathway in RHA rats reduced motor impulsivity, without affecting risk-related decision-making. Conversely, inhibition of the mPFC-NAc pathway had no effect in RLA rats. Our results suggest that the mPFC-NAc pathway controls motor impulsivity, but has limited involvement in risk-related decision-making. Our findings suggest that reducing fronto-striatal activity may help attenuate motor impulsivity in patients with impulse control dysregulation like ADHD or SUD.

## Introduction

Excessive impulsive behaviors, including motor impulsivity and risk-related decision making, are commonly associated with psychiatric disorders such as attention– deficit/hyperactivity disorder (ADHD), bipolar disorders, and substance use disorders^1–5^. Thus, exploring the neurobiological circuits underlying abnormal motor impulsivity and risk-related decision-making is crucial to bring new insights into the mechanisms contributing to mental health disorders.

Human studies have linked a reduced activity in the medial prefrontal cortex (mPFC) to both high motor impulsivity and riskier decision-making^6,7^. However, preclinical studies investigating mPFC alterations in relation to motor impulsivity have yielded mixed results. Excitotoxic lesions or reversible inactivation of the mPFC either increased^8–10^ or had no effect on motor impulsivity^11,12^, but impaired risk-related decision-making^11–13^. Despite these contrasting findings, lower cortical activity has long been studied as a neural basis for high levels of impulsive behaviors. In line with this, lower cortical levels of cFos expression, have been reported in poor compared to good decision-makers^14^. More recently, optogenetic or chemogenetic silencing of neuronal activity within specific subregions of the mPFC, including the anterior cingulate (ACC), prelimbic (PL) and infralimbic (IL) cortices, revealed that different mPFC regions might differentially, and even oppositely, control impulsive behaviors. Inhibition of ACC or IL reduced motor impulsivity^15,16^, while inhibition of PL increased it^15^. Although contrasting with pharmacological inactivation or lesions of the PL and IL, these studies all point towards opposite roles for the dorsal and ventral mPFC in impulsive behaviors. Few studies have investigated the specific cortical projections involved in impulse control, and further studies are still required to disentangle the cell- and circuit-specific involvement of the mPFC in motor and risk-related impulsivity. The mPFC sends widespread long-range projections across the brain^17,18^, and among the structures receiving input from the mPFC^17,18^, the nucleus accumbens (NAc) is of particular interest. Indeed, inhibition of the fast spiking GABAergic interneurons in the NAc increases motor impulsivity^19^ and low levels of cFos have been found in the NAc of poor vs. good decision-makers^14^. Besides, converging evidence indicates that both motor and risk-related impulsivities are linked to exaggerated levels of evoked dopamine (DA) release in the NAc^20,21,22^. Given that cortical glutamate inputs to the NAc can regulate accumbal DA^23–25^, alterations in cortical activity could potentially disrupt DA release in the NAc, leading to abnormal levels of motor and risk-related impulsivities.

Only a few studies have explored the role of the mPFC-NAc pathway on motor impulsivity and risk-related decision-making. Simultaneous excitotoxic lesions or pharmacological inactivations of both the mPFC and contralateral NAc have been used to induce widespread functional disconnections of these two regions, leading to increased motor impulsivity^26,27^ but no impairment in risk-related decision-making^28^. To our knowledge, only one study has investigated the role of neurons projecting from the ventral mPFC to the NAc, showing that chemogenetic activation of these neurons reduced motor impulsivity in a rat model of binge-like eating^29^. Yet, data are missing regarding the involvement of the mPFC-NAc projecting neurons on risk-related decision-making.

The Roman High-(RHA) and Roman Low Avoidance (RLA) sublines exhibit phenotypic differences in impulsivity, with RHA rats showing higher motor impulsivity^20,30–32^ and risk-related decision-making compared to RLA rats^30,31^. Hence, RHA and RLA rats offer an ideal model for studying the contribution of the mPFC in motor and risk-related impulsivity. Here, we investigate the impact of the chemogenetic modulation of the mPFC-NAc projecting neurons on impulsive behaviors in RHA and RLA rats. To this aim, we used an intersecting viral strategy to selectively activate the mPFC-NAc projecting neurons in RHA or inhibit them in RLA, and assessed the impact of these manipulations on motor and risk-related impulsivities using the rodent gambling task (rGT). Further, we used in vivo positron emission tomography (PET) imaging with [^18^F]Fluorodeoxyglucose ([^18^F]FDG) to examine brain glucose metabolism as an index of regional neuronal activity^33,34^, aiming to explore alterations in regional functional activity in relation to motor and risk-related impulsivity in both rat lines.

## Material and Methods

### Animals

Experiments were conducted in outbred male RHA and RLA rats from our colony at the University of Geneva. Rats were aged 2 months at the beginning of the experiment. Animals were housed in pairs on a 12:12 hours light-dark cycle (lights on at 7:00 am). Animals were food restricted to 85-90% of their free-feeding weight. Water was available ad libitum. Experiments were conducted under the Swiss Federal Law on animal care and were approved by the Animal Ethics Committee of the Geneva Canton.

### Timeline of the study

The timeline of the study is depicted in Figure 1. Rats were first trained in the rat Gambling Task (rGT). After at least 20 days of rGT training, rats underwent a stereotaxic surgery to be infused with viruses encoding the expression of chemogenetic Designer Receptors Exclusively Activated by Designer Drugs (DREADD), or mCherry alone (control condition), specifically in the mPFC-NAc projecting neurons. Following surgical recovery, rats were trained back in the rGT for a minimum of 20 sessions. Then, three rGT sessions were performed under saline treatment (i.p.) to establish baseline performance, followed by one session under the DREADD agonist clozapine-N-oxide (CNO, 1mg/kg). The effect of CNO vs. saline treatment was then assessed on locomotor activity tests performed at one-week interval and in a counterbalanced order. Finally, rats underwent two PET scans with [^18^F]FDG, one under saline and one under CNO treatment, one week-apart and in a counterbalanced order, to assess the effect of DREADD-mediated modulation of mPFC-NAc projections on brain metabolism.

**Figure 1.**
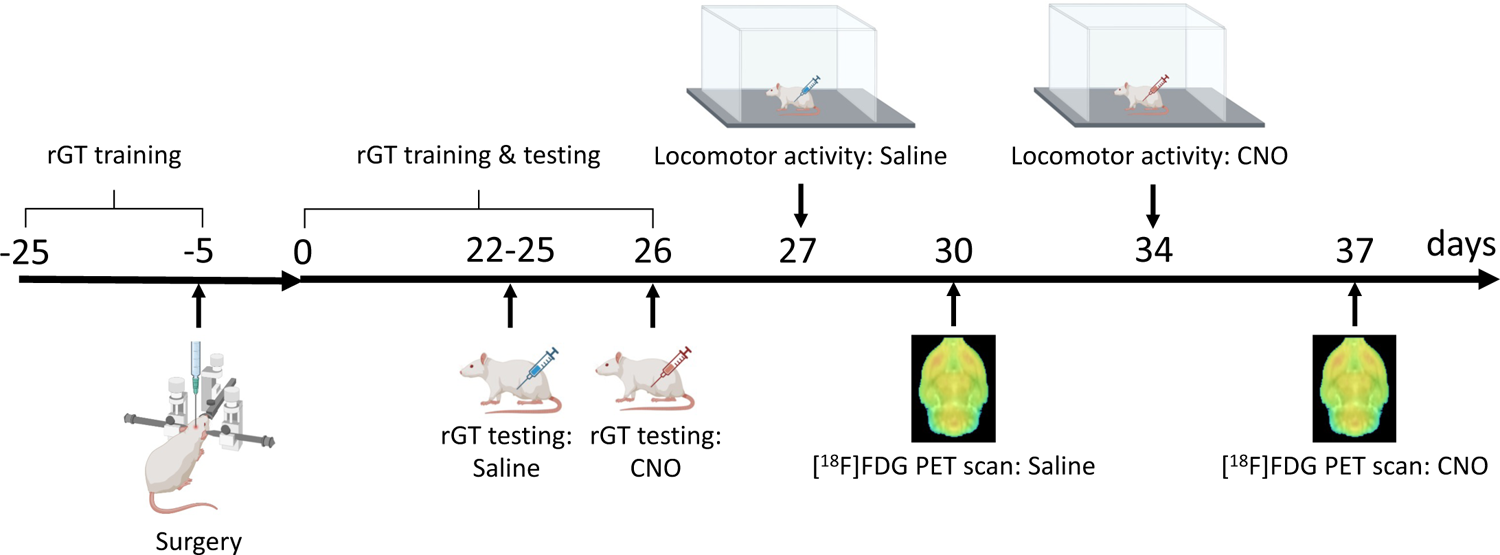
Experimental timeline of the study

### Drug

The exogenous DREADD ligand, CNO (HelloBio, Bristol, UK), was freshly prepared in 0.9% saline on the days of testing. All CNO injections were given i.p. at a dose of 1mg/kg, 30 minutes before testing. The dose of CNO was chosen based on previous studies^35,36^.

### Adeno-associated virus

Chemogenetic modulation of the mPFC-NAc pathway was achieved using a combined injection of two viral preparations. Animals received bilateral injections of a retrograde adeno-associated virus (AAV) encoding a Cre-recombinase AAV-retro/2-hSyn-EGFP-Cre (8.8×1012 vg/mL; Viral Vector Facility, Zurich, Switzerland) into the NAc. To target specifically the mPFC-NAc pathway, animals were injected bilaterally in the mPFC with an anterograde Cre-dependent AAV5-hSyn-DOI-hM3Dq-mCherry (7×10¹² vg/mL, plasmid #44361, Addgene) for RHA rats (hM3Dq-group, n=13), an anterograde Cre-dependent AAV5-hSyn-DOI-hM4Di-mCherry (7×10¹² vg/mL, plasmid #44362, Addgene) for RLA rats (hM4Di-group, n=14), or an anterograde Cre-dependent AAV5-hSyn-DOI-mCherry (7×10¹² vg/mL, plasmid #50459, Addgene) for control groups (mCherry control groups; n=12 RHA; n=7 RLA).

### Surgeries

Surgical procedure details are provided in the Supplementary materials. Briefly, animals were anesthetized placed in a stereotaxic apparatus (RWD Life Science, Mainz, Germany). Viral preparations were injected in a volume of 500nL per site, at 3nL/sec using a Nanoject III injector (Drummond Scientific Company, Broomal, PA, USA). Viruses were injected in the NAc (AP=1.07; ML=±0.12; DV=-0.63 from dura) or the mPFC (AP=1.2; ML=±0.04; DV=-0.37 from dura). Coordinates AP and ML are expressed in cm from the *lambda*. Then, the scalp was sutured and rats returned to their home cage for post-operative care. Animals were allowed to recover for 1 week, during which food and water were provided ad libitum.

### Rat gambling task

Training and testing in the rGT were performed as previously described^30,37^. Additional details about the procedure are provided in the Supplementary materials. Briefly, rats were trained to nosepoke into four holes (P1, P2, P3, P4), after an inter-trial interval (ITI), to receive food pellet rewards. Each hole was associated with different probabilities of receiving a different amount of reward or a time-out (TO) punishment of varying duration (Table 1). When a trial was punished, the chosen hole blinked at 0.5 Hz for the duration of the TO. Choosing the holes associated to P1 and P2 (i.e., optimal choices) throughout the session resulted in more pellets earned at the end of the session, compared to the P3 and P4 options (i.e., non-optimal choices), leading to longer TO punishment and less pellets earned. Risk-related decision-making was indexed with the choice score (%optimal choices - %non-optimal choices). Responses during the 5sec-ITI were scored as premature responses, and signaled by a 5sec TO period. The percentage of premature responses [(#premature responses/#trial initiated)*100] was an index of motor impulsivity. After 20 sessions of rGT, saline was injected i.p. 30min before sessions to control for the stress of injections, for a minimum of 3 sessions. A mixed factorial analysis of variance (ANOVA) was performed on 3 selected days for baseline performance, with line (i.e. RHA or RLA) as between-line factor, and choice score over the 3 selected days as within-subject factor. The 3 days selected for baseline were considered stable if no main effect of session nor line x session interaction were observed. When stable baseline was observed, the effect of mPFC-NAc pathway modulation on motor impulsivity and risk-related decision-making was assessed in a single-session of rGT performed after the administration of CNO.

**Table 1:**
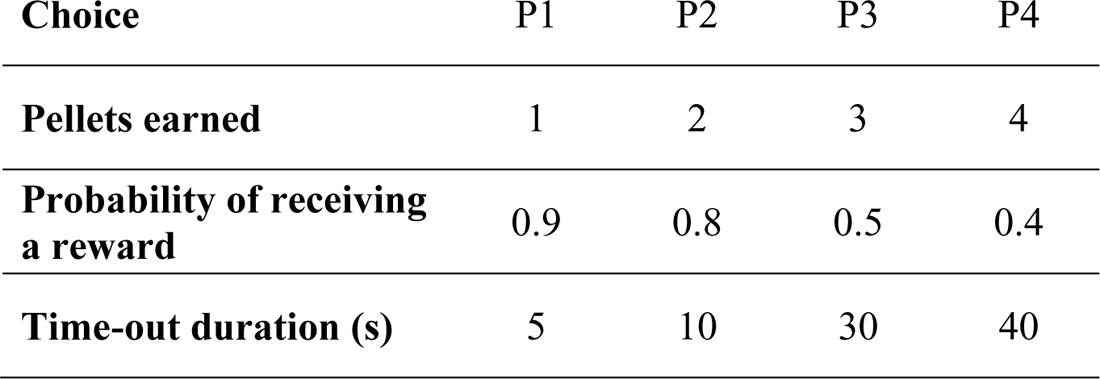
Choice contingencies in the rGT (adapted from ^37^)

### Locomotion

Locomotor activity was assessed in open-fields (Actimot, TSE Systems, Bad Homburg, Germany), equipped with two rows of 16 photocell beams to measure horizontal activity. Light was dimmed at approximately 10 lux intensity. Rats received an i.p. injection of CNO or saline before being placed in the center of the open-field. A second test was performed one week later, with counterbalanced treatment. Locomotion was monitored for 60min, and total distance traveled was averaged in 10-min blocks.

### [^18^F]FDG PET scanning and analyses

Rats underwent two PET scans with [^18^F]FDG using a Triumph II scanner (Trifoil Imaging, Northridge, CA, USA). Rats were fasted overnight before each [^18^F]FDG PET scan, to avoid competition between endogenous glucose uptake and [^18^F]FDG uptake^38,39^. On the day of the scan, animals received an i.p. injection of saline or CNO 30min prior the injection of [^18^F]FDG. Animals were injected i.v. with 27.7±3.1 MBq [^18^F]FDG (sourced from the University Hospitals of Geneva, Switzerland) for RHA rats, and 28.1±3.3 MBq [^18^F]FDG for RLA rats. Rats were then returned in their home cage where they were kept warm using an infrared lamp for the next 45min^40^ (i.e., uptake period). Two rats per scan were anesthetized and positioned head-to-head in a custom-made bed in the PET gantry. Fifty minutes after receiving [^18^F]FDG, a 20-min emission scan was performed. Dynamic PET images were acquired in list mode and reconstructed into 4 time-frames (300s duration each) using the ordered subsets expectation maximization algorithm and 20 iterations.

Image analyses were performed using PMOD software (version 4.0, PMOD Technologies Ltd., Zurich, Switzerland). Individual dynamic PET images were averaged over the 20-min of data acquisition to create PET summation images. [^18^F]FDG brain templates were then generated for RHA and for RLA rats using a procedure previously described^41^. Briefly, PET summation images of 10 RHA rats were cropped to brain only images. From these images, one representative summation PET image was selected as the reference. Each individual summation scan was then spatially normalized to the reference image using an automated elastic transformation. Subsequently, all normalized images were averaged to generate an average image of the RHA rat brain. A mirrored version of this image was created by flipping the average image of the RHA rat brain from left to right. These two RHA brain images were then averaged into a final RHA brain [^18^F]FDG template. The same procedure was applied to the PET summation images of 10 RLA rats to generate a RLA brain [^18^F]FDG template.

Individual PET images were automatically co-registered to the corresponding RHA or RLA brain template using rigid registration based on normalized mutual information. When necessary, manual micro-adjustments were performed to align individual PET images to the corresponding template. Dynamic PET images were converted into Standardized Uptake Value (SUV) according to the formula [C/(D_inj_/W)], where C is the radioactivity concentration (kBq/cc), D_inj_ is the injected dose of [^18^F]FDG (kBq), and W is the weight of the animal (g). Then, a region of interest (ROI) template adapted from the ROI template of Schiffer and colleagues^40^ was created including the following regions: mPFC, NAc, dorsal striatum (DST), orbitofrontal cortex (OFC), thalamus (THAL). ROIs consisted of circular ROIs positioned bilaterally and on the central planes of each structure to minimize partial voluming. The ROI template was applied to the individual dynamic PET images and [^18^F]FDG SUV were extracted for each ROI. For each animal, the SUV were averaged over the 4 time-frames to obtain a final SUV for each region (SUV_ROI_). As previously described^42^, SUV_ROI_ were then normalized using a whole-brain normalization factor (average whole-brain uptake of all animals in one group/individual whole-brain uptake). These normalized SUV (NormSUV) were then used for analyses.

### Tissue preparation and Immunohistochemistry

Additional details are provided in Supplementary materials. Briefly, rats were transcardially perfused with 4% paraformaldehyde (PFA, Sigma-Aldrich, Steinhem, Germany), Brains were extracted, frozen and cut into 40 μm slices, Free-floating slices were stained with Hoescht (Abcam, #ab828550). Fluorescent images were acquired using a widefield fluorescence slide scanner microscope (Zeiss Axioscan Z1, Gottingen, Germany) and analyzed using Zen 2.0 software (Zeiss, Gottingen, Germany).

### Statistical analyses

Normality of the data was assessed using a Shapiro-Wilk test. At baseline (saline condition), between lines differences in the % of premature responses, choice scores, % of omissions and total trials during the rGT were assessed using a Student’s t-test for normal data or a Mann-Whitney U-test for non-normal data. Prior to ANOVA analyses, homogeneity of variances was verified with Levene’s test. Sphericity was assessed using Mauchly’s test, and the degree of freedom were corrected using Greenhouse-Geisser correction if needed. Between-lines differences in regional [^18^F]FDG NormSUV values under saline condition were assessed using a mixed factorial ANOVA with lines as between-subject factor and brain region as within-subject factor. Significant effects were analyzed with post-hoc Student’s t-tests, where appropriate. Within lines, the effect of mPFC-NAc pathway modulation on rGT performances, locomotor activity and [^18^F]FDG NormSUV values was assessed using a mixed factorial ANOVAs with virus (DREADD vs. Control) as between-subject factor, and treatment (CNO vs. Saline) as within-subject factor. Data are presented as mean ± SEM, and considered significant when p<0.05. Statistical analyses were performed with Graphpad Prism (Graphpad Software 9.0.2, San Diego, CA, USA).

## Results

### Baseline performance in the rGT

Consistent with previous data^30,31^, RHA rats displayed a higher percentage of premature responses at baseline (t=4.23, df=44, p<0.001, Fig.2A), indicating higher levels of motor impulsivity compared to RLA rats. RHA rats also tended to choose the riskiest options (i.e., P3 and P4) more often than RLA rats, as indicated by a nearly significant lower choice score in RHA compared to RLA rats (U=179, p=0.07; Fig. 2B). The percentage of omissions was lower in RHA than in RLA rats (t=2.68, df=44, p<0.05, Fig. 2C), although both rat lines performed a similar number of trials in the task (t=1.02, df=44, p=0.31, Fig. 2D). Finally, the percentages of premature responses were negatively correlated with the choice scores (r=-0.52, p<0.001; Fig.2E), indicating a positive correlation between motor impulsivity and risk-related decision-making.

**Figure 2:**
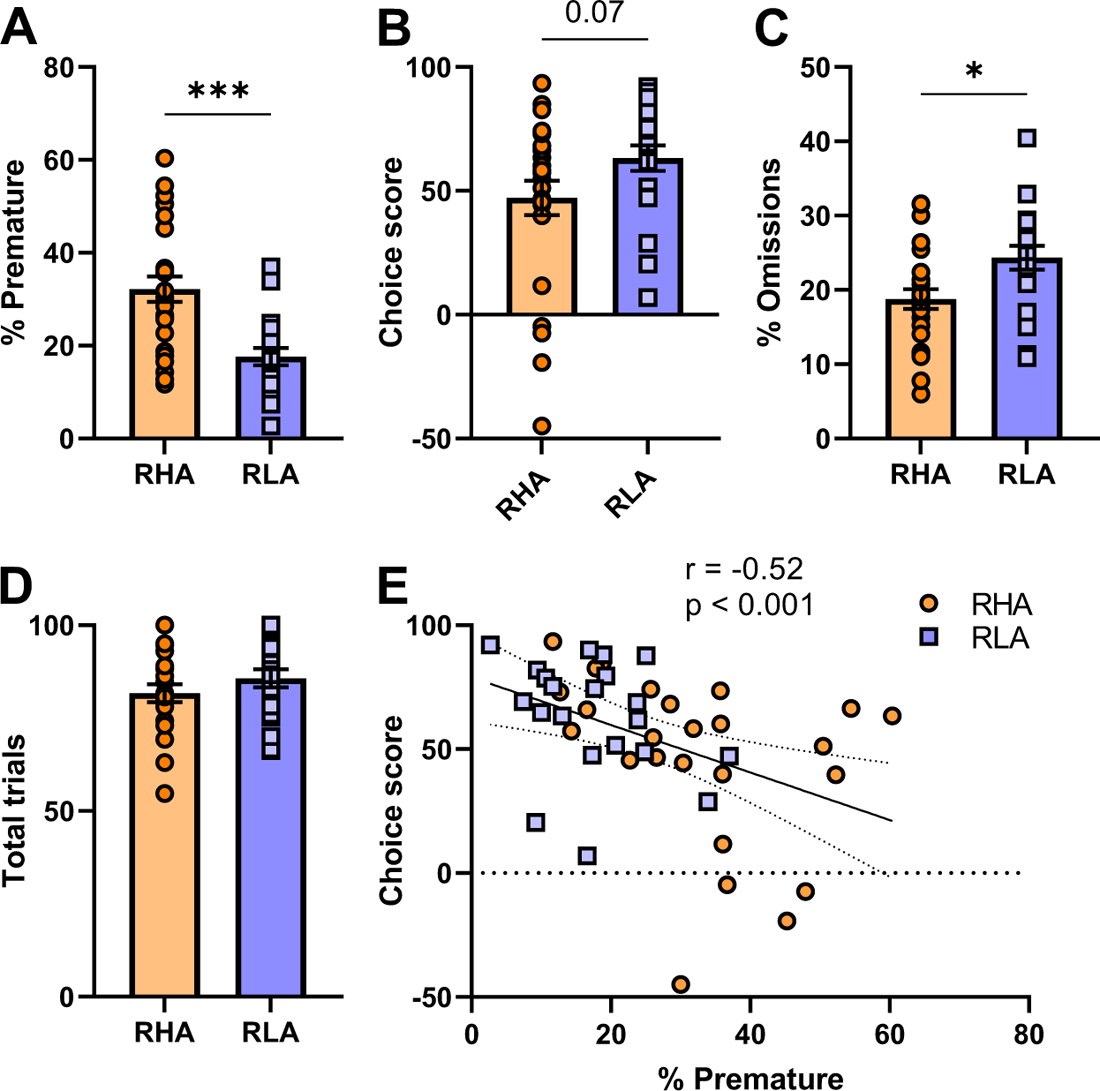
Baseline performances in the rGT. Compared to RLA rats, RHA rats exhibited **(A)** a higher percentage of premature responses, **(B)** tended to have a lower choice score, **(C)** made fewer omissions but **(D)** completed a similar number of total trials. **(E)** Choice scores and percentages of premature responding were negatively correlated in the rGT. Data are represented as mean ± SEM; *p<0.05; ***p<0.001

### Modulation of mPFC-NAc pathway on rGT performances

Representative images of viral expression in the mPFC are depicted in Fig. 3B. Widefield microscope images show GFP-tagged neurons indicating Cre-recombinase expression (left, GFP), and mCherry-tagged neurons indicating Cre-dependent viral expression (middle, mCherry) restricted to GFP-expressing neurons (right, merge). In RHA rats, a mixed factorial ANOVA on premature responding (Fig. 3C) revealed no main effect of virus (F_1,23_=0.29, p=0.60) but a main effect of treatment (F_1,23_=10.11, p=0.0042) and a virus x treatment interaction (F_1,23_=6.24, p=0.020). Post-hoc comparisons indicated that, relative to saline, CNO induced a significant reduction of premature responding in the RHA-hM3Dq group (p=0.0009), but not in the RHA-mCherry control group (p=0.87). On the other hand, there was no main effect of treatment (F_1,19_=0.023, p=0.88) or virus (F_1,19_=0.38, p=0.55), and no treatment by virus interaction (F_1,19_=0.083, p=0.78) on premature responding in RLA rats (Fig. 3C). Regarding the choice score (Fig. 3D), CNO had no effect in either RHA (treatment: F_1,23_=1.17, p=0.29; virus: F_1,23_=0.029, p=0.87; treatment x virus: F_1,23_=0.060, p=0.81) or RLA rats (treatment: F_1,19_=0.86, p=0.37; virus: F_1,19_=3.13, p=0.09; treatment x virus: F_1,19_=1.40, p=0.25). The effect of CNO on other parameters of the rGT are presented in Fig. S1. In RHA rats, a mixed factorial ANOVA on the percentage of omissions revealed a main effect of treatment (F_1,23_=6.22, p=0.02), but no main effect of virus (F_1,23_=0.19, p=0.66) or treatment x virus interaction (F_1,23_=0.81, p=0.38; Fig. S1A), indicating that CNO increased the percentage of omissions in both RHA-hM3Dq and RHA-mCherry groups. CNO had no effect on omissions in RLA rats (treatment: F_1,19_=0.87, p=0.36; virus: F_1,19_=0.25, p=0.62; treatment x virus F_1,19_=0.22, p=0.65; Fig. S1B). Finally, we observed no effect of CNO on the total number of trials in RHA (treatment: F_1,23_=0.71, p=0.41; virus: F_1,23_=0.10, p=0.76; treatment x virus: F_1,23_=0.15, p=0.70; Fig. S1C) or in RLA rats (treatment: F_1,19_=1.84, p=0.19; virus: F_1,19_=0.28, p=0.28; treatment x virus: F_1,19_=0.02, p=0.89; Fig. S1D). Collectively, these results indicate that activation of the mPFC-NAc projecting neurons in RHA reduced their motor impulsivity, while inhibition of this pathway had no effect in RLA rats. Additionally, our findings suggest that the mPFC-NAc pathway likely contribute to motor impulsivity, but have a limited role in risk-related impulsivity.

**Figure 3:**
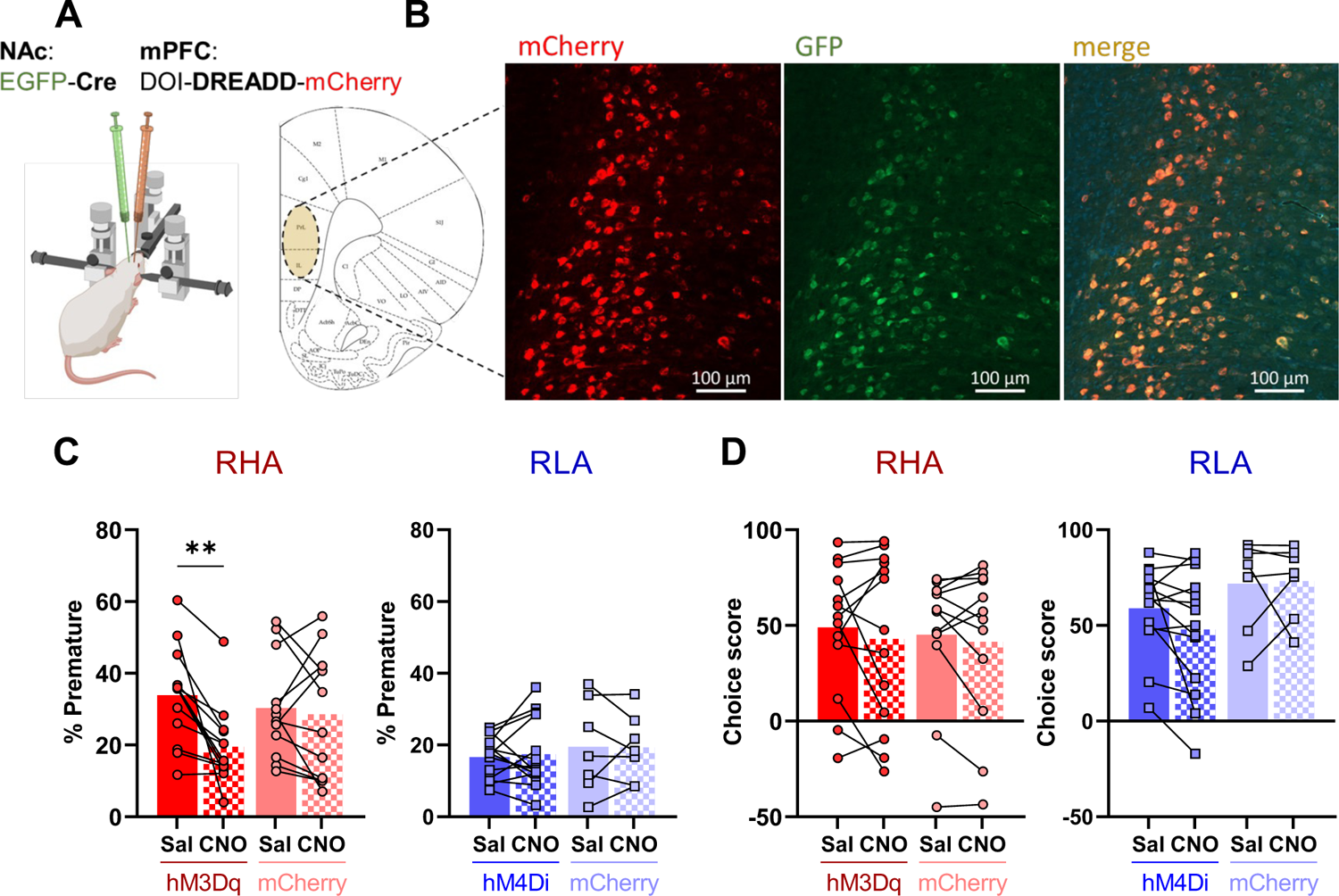
Effect of modulation of the mPFC-NAc projections on rGT performances. **(A)** Each rats received bilateral injections of a retrograde Cre-GFP virus into the NAc, along with bilateral injections of a Cre-dependent DREADD virus into the mPFC as follows: hM3Dq-mCherry virus for the RHA-hM3Dq group, hM4Di-mCherry virus for the RLA-hM4Di group and mCherry virus for RHA-control and RLA-control groups. **(B)** Representative images showing Cre-expressing neurons in green (left panel), mCherry-tagged neurons indicating Cre-conditional viral expression (middle panel) and co-localization of both GFP and mCherry in the mPFC (right panel). **(C)** Relative to saline, CNO-induced activation of mPFC-NAc projections reduced premature responding in the RHA-hM3Dq group but not in the RHA-mCherry group, whereas CNO-induced inhibition of mPFC-NAc projections had no effect in RLA-hM4Di or in RLA-mCherry groups. **(D)** When compared to saline, CNO treatment had no effect on choice scores in any of the RHA or RLA groups. **p<0.01

### Locomotor activity

To assess whether the observed CNO-induced effects on rGT performance were influenced by changes in general locomotion, we examined the locomotor activity of both RHA and RLA rats following saline and CNO treatment (Fig. 4). In RHA rats (Fig.4A), a mixed factorial ANOVA revealed a main effect of treatment (F_1,23_=5.74 p=0.025), but no main effect of virus (F_1,23_=2.78, p=0.11) and a trend for a treatment x virus interaction (F_1,23_=3.99, p=0.058). Although the interaction effect did not reach conventional statistical significance, we conducted post-hoc comparisons, which revealed that activation of the mPFC-NAc pathway increased locomotor activity in the RHA-hM3Dq group (p<0.01) but not in the RHA-mCherry control group (p=0.95). In RLA rats (Fig. 4B), a mixed factorial ANOVA revealed a main effect of treatment (F_1,19_=5.61, p=0.029), no main effect of virus (F_1,19_=2.80, p=0.11) and a trend for a treatment x virus interaction (F_1,19_=4.15, p=0.056). Post-hoc comparisons indicated that inhibition of the mPFC-NAc pathway reduced locomotor activity specifically in the RLA-hM4Di (p<0.01) but not in the RLA-mCherry control group (p=0.97). These data suggest that the CNO-induced reduction in motor impulsivity observed in RHA rats during the rGT is not attributable to a reduction in their locomotor activity.

**Figure 4:**
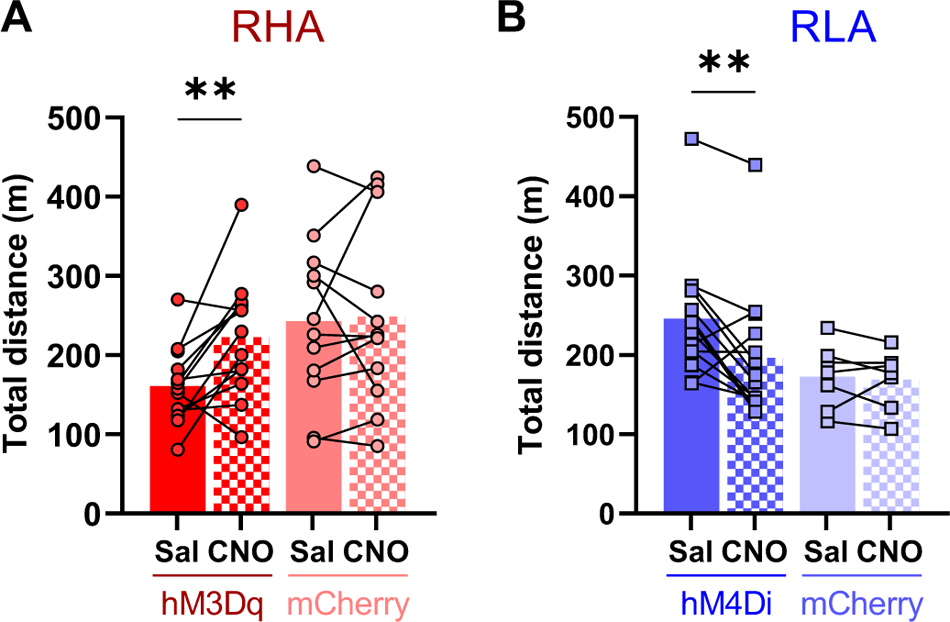
Effect of modulation of the mPFC-NAc projections on locomotor activity. **(A)** In RHA rats, CNO increased locomotor activity in the RHA-hM3Dq but not in the RHA-mCherry control group, when compared to saline. **(B)** In RLA rats, CNO decreased locomotor activity in the RLA-hM4Di but not in RLA-mCherry control group. **p<0.01

### Baseline differences and CNO effects on regional [^18^F]FDG uptake in RHA and RLA rats

Mean parametric maps of [^18^F]FDG NormSUV values comparing RHA and RLA rats at baseline (e.g. saline condition) are shown in Fig 5A. Under baseline saline conditions (Fig. 5B), a mixed factorial ANOVA revealed a main effect of brain region (F_3,121_=125.9, p<0.001), line (F_1,40_=22.68, p<0.001) and brain region x line interaction (F_5,200_=5.75, p<0.001) on [^18^F]FDG NormSUV values. Planned comparisons revealed that [^18^F]FDG NormSUV was lower in RHA compared to RLA rats in the mPFC (t=2.38, df=40; p<0.05), NAc (t=2.92, df=40; p<0.01), OFC (t=5.68, df=40; p<0.001) and THAL (t=9.27, df=40; p<0.001).

**Figure 5:**
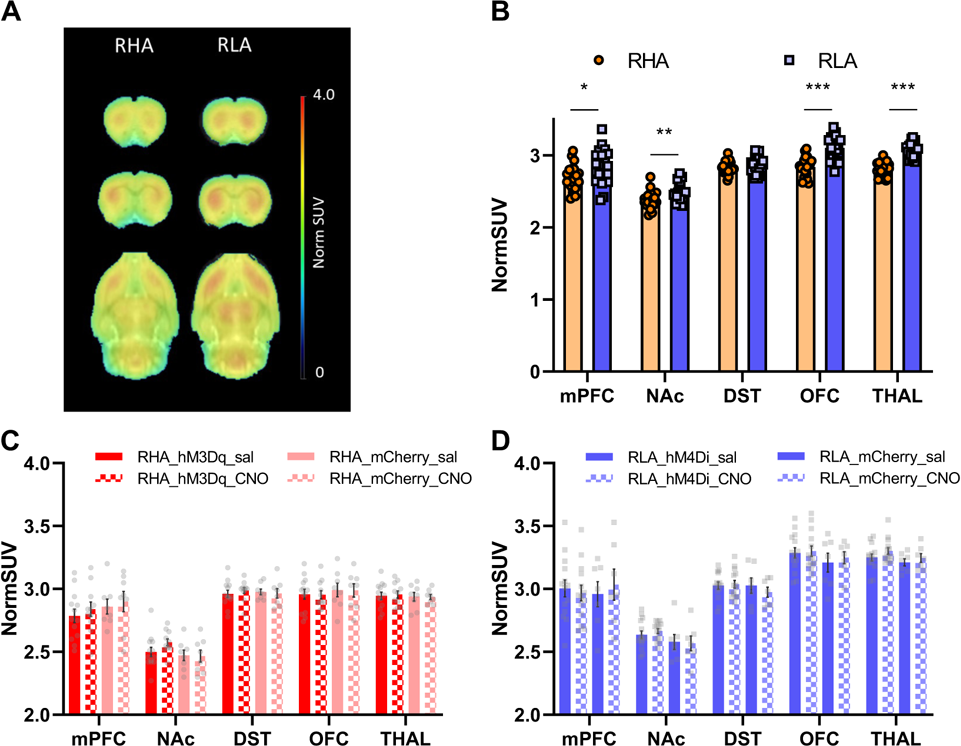
Effect of the modulation of the mPFC-NAc projections on [^18^F]FDG brain uptake. **(A)** Mean parametric maps of [^18^F]FDG NormSUV values in RHA (left) and RLA rats (right) are shown in coronal planes at the level of the mPFC (top), and striatum (middle) and in a horizontal plane (bottom). **(B)** [^18^F]FDG NormSUV values were lower in RHA rats (n=21) vs. RLA rats (n=21) in mPFC, NAc, OFC, and THAL, but did not differ between lines in DST and CING. **(C)** Compared to saline, CNO had no effect on [^18^F]FDG uptake in RHA-hM3Dq or RHA-mCherry groups or **(D)** between RLA-hM4Di and RLA-mCherry groups. Data are represented as mean ± SEM; *p<0.05; **p<0.01; ***p<0.001

The effects of CNO vs. saline vehicle on regional [^18^F]FDG NormSUV values are shown in Fig. 5C-D. In RHA rats (Fig. 5C), a mixed factorial ANOVA indicated no main effects of treatment or virus, and no treatment x virus interaction in the mPFC (treatment: F_1,17_=1.06, p=0.32; virus: F_1,19_=0.85, p=0.37; treatment x virus: F_1;17_=0.0009, p=0.98), NAc (treatment: F_1,17_=0.84, p=0.37; virus: F_1,19_=3.08, p=0.10; treatment x virus: F_1;17_=1.17, p=0.30), DST (treatment: F_1,17_=0.02, p=0.89; virus: F_1,19_=0.001, p=0.97; treatment x virus: F_1;17_=0.44, p=0.51), OFC (treatment: F_1,17_=0.17, p=0.69; virus: F_1,19_=0.52, p=0.48; treatment x virus: F_1;17_=0.09, p=0.76), or THAL (treatment: F_1,17_=0.0003, p=0.99; virus: F_1,19_=0.08, p=0.78; treatment x virus: F_1;17_=0.04, p=0.84). Similarly, in RLA rats (Fig. 5D), there were no main effects of treatment or virus, and no treatment x virus interaction in the mPFC (treatment: F_1,19_=0.15, p=0.70; virus: F_1,19_=0.01, p=0.92; treatment x virus: F_1;19_=1.28, p=0.27), NAc (treatment: F_1,19_=0.14, p=0.71; virus: F_1,19_=2.46, p=0.13; treatment x virus: F_1;19_=0.71, p=0.41), DST (treatment: F_1,19_=0.62, p=0.44; virus: F_1,19_=0.43, p=0.52; treatment x virus: F_1;19_=1.32, p=0.26), OFC (treatment: F_1,19_=0.55, p=0.47; virus: F_1,19_=1.04, p=0.32; treatment x virus: F_1;19_=0.13, p=0.72), or THAL (treatment: F_1,19_=1.99, p=0.17; virus: F_1,19_=2.17, p=0.16; treatment x virus: F_1;19_=0.09, p=0.77). Collectively, these findings suggest that no detectable effect of CNO was observed on [^18^F]FDG NormSUV values in any regions.

## Discussion

To our knowledge, this is the first study to investigate the role of mPFC-NAc projecting neurons in both motor impulsivity and risk-related decision-making using a within-subject design. Our findings showed that high impulsive RHA rats display lower cortical functional activity compared to low impulsive RLA rats. Furthermore, we showed that chemogenetic stimulation of mPFC-NAc projecting neurons decreased motor impulsivity in high-impulsive animals, whereas chemogenetic inhibition of this pathway had no effect in low-impulsive animals. Conversely, chemogenetic modulation of the cortico-striatal projecting neurons had no effect on risk-related decision-making, regardless of baseline levels of impulsivity. Our results suggest that the mPFC-NAc projecting neurons are involved in motor impulsivity but have limited impact on risk-related impulsivity. Therefore, despite being positively correlated^30,31,43^, motor and risk-related impulsivities are likely controlled by distinct neuronal circuits.

This is the first study to investigate differences in brain metabolism using [^18^F]FDG and PET imaging in high vs. low impulsive rats. We observed that RHA rats exhibited lower [^18^F]FDG uptake in brain regions that have been associated with impulsive behaviors, such as the mPFC^26,44^, NAc^19^, OFC^45^ and thalamus^46^. This is consistent with clinical data showing lower cortical glucose metabolism in patients with ADHD^47^ or borderline personality disorders^48–50^, who also display low impulse control^51^. Our findings suggest a potential alteration of the activity of the cortico-striato-thalamic circuit in RHA compared to RLA rats. Alteration of the activity of this circuit has been reported in humans in patients with schizophrenia^52^, a disorder that has repeatedly been associated with high impulsivity^53^. Our data bring new insight on the potential underpinnings of high motor impulsivity, and suggest that alterations in brain metabolic activity, especially in the cortico-striato-thalamic circuit, may be associated to elevated levels of impulsive behaviors.

Chemogenetic activation of mPFC-NAc projecting neurons decreased motor impulsivity in high impulsive animals, suggesting that a hypoactivity of this pathway contributes to this facet of impulsivity. Notably, our observation of increased locomotion upon activating the mPFC-NAc neurons indicates that the decrease in motor impulsivity reported here unlikely results from a CNO-induced change in locomotor function. Besides, our results are in line with studies showing that non-specific pharmacological lesions or inactivation in the mPFC increased motor impulsivity^8–10^. Additionally, our findings are also congruent with previous studies reporting increased motor impulsivity following optogenetic inhibition of cortical layer 5 of the mPFC^54,55^, the main cortical output layer. Here, we expanded upon these prior findings by presenting new insights into the neural circuit underlying motor impulsivity, highlighting the significant contribution of mPFC neurons projecting to the NAc. The cortical neurons projecting to the NAc are mainly glutamatergic and, interestingly, high impulsive animals display lower glutamate in the NAc compared to low impulsive rats^56^, suggesting that disrupted glutamatergic transmission between mPFC and NAc is involved, at least partly, for high impulsive behaviors.

In contrast to the decrease in motor impulsivity observed with chemogenetic activation of mPFC-NAc projecting neurons in high-impulsive rats, chemogenetic inhibition of these neurons had no impact on motor impulsivity in low-impulsive rats, indicating that inhibition of this pathway is not sufficient to induce high motor impulsivity. One possibility to explain this result may be that high impulsive behaviors observed in RHA rats are due to a combination of several factors. For instance, in addition to lower cortical activity, RHA rats exhibit abnormalities in meso-striatal DA signaling, such as heightened evoked DA release in the striatum and lower densities of DA D_2/3_ receptors in the striatum, relative to RLA rats^20,57^, both of which have been related to motor impulsivity^20,21^. Interestingly DA acts as a neuromodulator in the NAc, and regulates the cortical glutamatergic inputs^58,59^. Thus, the activity of the NAc neurons may depend not only on cortical afferences, but also on DA release. Therefore, high motor impulsivity in RHA rats may result from a combined lower activity of the mPFC-NAc pathway along with elevated DA release, compared to RLA rats. Consequently, inhibition of the mPFC-NAc pathway alone may not be sufficient to induce motor impulsivity in RLA rats. Further studies using combined inhibition of the cortico-striatal pathway along with activation of the DA terminals in the striatum may be necessary to increase motor impulsivity in RLA rats.

We observed no effect of the modulation of mPFC-NAc pathway on risk-related decision-making. Although studies on the involvement of this pathway on risky decision-making are scarce, our results contrast with previous findings showing an involvement of the mPFC^13,60^ on one hand, and the NAc^61^ on the other hand in risk-related impulsivity. However, these studies suggested that the ventral part of the mPFC (IL) and shell part of the NAc (NAcSh) are involved in sensitivity to punishment (e.g., reward omission) while the dorsal part of the mPFC (PL) and core part of the NAc (NAcC) are involved in updating choice values. During the rGT, the probabilities of receiving rewards or punishments do not change across sessions, suggesting that IL may be more involved than PL in this task. This hypothesis is in line with data suggesting an opposite role for the dorsal and ventral part of the mPFC in risky decision-making^62^. In our study, we targeted the mPFC neurons projecting to the NAc, including both the PL and IL cortical subregions projecting onto the NacC and NAcSh, respectively^18^, which may mask the respective contribution of each region to risk-related decision-making. A combined but opposite activity of these subregions may then contribute to optimal choices in the rGT, and modulation of even more specific circuits may be required to evidence the involvement of PL and IL in risk-related decision-making.

Given the positive correlation between motor impulsivity and risk-related decision-making reported here and previously by others^30,31,43^, the selective involvement of the mPFC-NAc projecting neurons in one facet of impulsivity but not the other is surprising. However, such a selective effect has been previously observed following pharmacological inactivation of different cortical regions, which altered decision-making without impacting motor impulsivity^11,12^. This suggests that, while correlated, these two facets of impulsivity likely share neurobiological substrates but involve at least partly distinct pathways. As such, other brain pathways could underlie risk-related decision-making, such as the neurons projecting from the basolateral amygdala (BLA) to the NAc^63,64^. As the mPFC sends projections to NAc-projecting BLA neurons^65^, risk-related decision-making may involve the activity of mPFC-BLA-NAc circuit, while motor impulsivity may be influenced by more direct mPFC-to-NAc projections. Besides exploring the involvement of mPFC-NAc projecting neurons in motor and risk-related impulsivity, our study aimed to assess whether chemogenetic modulation of this pathway was detectable using PET [^18^F]FDG. Unfortunately, in our experimental conditions, CNO-induced changes in neuronal activity were not detected using [^18^F]FDG PET. These results were unexpected considering that a recent research was able to detect changes in glucose metabolism following chemogenetic modulation of the nigro-striatal projecting neurons^42^. However, this latter pathway may be denser than the mPFC-to-NAc pathway, and may therefore induce more drastic metabolic changes that can be detected with [^18^F]FDG PET. Thus, our results indicate that while chemogenetic manipulation of the mPFC-NAc neurons is sufficient to induce behavioral changes, its impact on glucose metabolic activity in mPFC may fall below the detection threshold of [^18^F]FDG PET imaging.

Interestingly, lower cortical activity has been associated with addiction. Indeed, neuroimaging studies in addicted patients have reported lower activity of the frontal cortex^7,66^, which has been associated with higher levels of motor impulsivity^67^. In addition to their low cortical metabolism observed here, high motor impulsive RHA rats exhibited greater cocaine intake^21,30,68^, higher levels of drug-primed relapse^30^ and lower gray matter volume compared to their low impulsive RLA counterparts^69^. Interestingly, mPFC activity has been implicated in several characteristics of addictive-like behaviors. Inhibition of mPFC accelerated acquisition of cocaine-taking^70^, and lesioning or inactivation of PL decreased cocaine seeking, but also stress-, cue-, context- or cocaine-induced reinstatement^71^, while inhibition or inactivation of IL increased drug-induced reinstatement^72^, and functional disconnection between IL and NAcS induced reinstatement of drug-seeking following extinction^73,74^. Together with our results, this suggests that hypoactivity in the fronto-striatal neurons may serve as a marker for both high motor impulsivity and vulnerability to drug consumption.

While our study focused on male rats, there are gender differences in impulsivity and risk-related decision-making, with female rats displaying lower levels of both motor^75^ and risk-related^76,77^ impulsivities compared to males. Furthermore, cortical activity has been associated with motor impulsivity in male, but not in female rats^78^. In clinical studies, mixed results have been reported. During a task measuring motor impulsivity, some have found a higher^79,80^, while others reported lower cortical activity in men compared to women^81^. Thus, future studies should consider gender differences to further elucidate how gender influences the neural underpinnings of motor impulsivity and risk-taking.

In conclusion, our study showed that high impulsive RHA rats exhibit lower cortical metabolism than low-impulsive RLA rats, suggesting a functional cortical hypoactivity in the former strain. PET investigation of brain glucose metabolism suggested the involvement of an extended neural circuit, encompassing cortical areas, ventral striatum and thalamus, in impulsive behaviors. In particular, our findings shed light on the role of the mPFC-NAc projecting neurons in motor impulsivity and risk-related decision-making. Activation of these neurons effectively reduced motor impulsivity in high-impulsive RHA rats, emphasizing the involvement of a reduced cortical top-down inhibitory control on the NAc in this facet of impulsivity. However, the absence of effect on risk-related decision-making despite a positive correlation between motor and risk-related impulsivity^30,31,43^, highlights that these two impulsivity facets are underpinned by different, yet likely overlapping, neural circuits. Although further studies are still required to offer a more refined understanding of the neural mechanisms involved in risk-related impulsivity, our study provides a better understanding of the neurobiological substrates underlying high motor impulsivity. Importantly, our results suggest that dampening cortico-striatal activity may be a potential strategy to attenuate excessive motor impulsivity in patients with impulse control dysregulation such as ADHD^3,4^ or SUD^2,5^.

## Acknowledgements

We thank Stéphane Germain for his assistance during the PET scans, and Steven Lang for his technical assistance in conducting part of the experiments. xThis study was funded by the Swiss National Science Foundation (SNSF; Grant Number: 31003A_179373). SNSF had no further role in study design; collection, analysis, or interpretation of data; writing of the report; or the decision to submit the manuscript for publication.

