## Supplementary materials for "Activation of the mPFC-NAc pathway reduces motor impulsivity but does not affect risk-related decision-making in innately high-impulsive rats"

**Supplementary Methods**

### Surgeries

Animals were anesthetized with isoflurane 3% in O_2_ and placed in a stereotaxic apparatus (RWD Life Science, Mainz, Germany). Anesthesia was maintained at 1.5-2% isoflurane in O_2_. Breathing, reflexes and body temperature were monitored throughout the surgery. An incision was made following the midline of the scalp, the skull was exposed and burr holes were drilled above the targeted coordinates. Viral preparations were injected in a volume of 500nL per site, at 3nL/sec using a Nanoject III injector (Drummond Scientific Company, Broomall, PA, USA). Viruses were injected in the NAc (AP=1.07; ML=±0.12; DV=-0.63 from dura) or the mPFC (AP=1.2; ML=±0.04; DV=-0.37 from dura). Coordinates AP and ML are expressed in cm from the *lambda*. After infusion, the glass capillary was left in place for 10 minutes to allow for diffusion of the viruses. After slow (i.e., 1min) removal of the capillary, the scalp was sutured and rats returned to their home cage for post-operative care. Buprenorphine (0.1 mg/kg) was provided s.c. 20 min before surgery, and every 6h during 48h post-surgery with a relay in the drinking water at night. Animals were allowed to recover for 1 week, during which food and water were provided ad libitum.

### Rat gambling task

Sessions took place in operant conditioning chambers (Med Associates Inc., St Albans, VT, USA), enclosed in sound-attenuated cubicles. Each chamber was composed of a food tray where Rodent Dustless Precision Pellets® (45 mg Noyes dustless pellets, TestDiet®, St Louis, MO, USA) were dispensed. On the opposite wall, 5 holes were available, positioned at 2.5cm above the floor of the box, equipped with a cue-light and infrared detector. The middle hole was not used during the rGT training and testing. The rGT was performed as originally described (Zeeb et al., 2009). Briefly, rats were first habituated to the operant boxes during 2 daily 30-min sessions. Then, rats were trained to nose-poke in each individual hole in daily 30-min sessions, during which responding in one illuminated hole was rewarded by a food reward (pellet). The order of the illuminated holes was pseudo-randomized. Animals were then trained in forced-choice sessions of rGT, during which one hole per trial was illuminated. After 3 forced-choice sessions, animals went through stereotactic surgery for intracranial injections of the viral preparations. One week after surgery, animals were trained back in 3 to 5 forced-choice sessions of rGT and were then tested in free-choice sessions of rGT. A trial started with a nose-poke in the food tray. After 5 seconds of inter-trial interval (ITI), the 4 holes were illuminated for 10 s, during which time rats had to choose which hole to nose-poke into. Each hole was associated with different probabilities P1 (0.9), P2 (0.8), P3 (0.5) and P4 (0.4) of receiving a different amount of reward (1, 2, 3, or 4 pellets, respectively), or a time-out (TO) punishment of varying duration (5, 10, 30, 40 seconds, respectively). When a trial was punished, the hole chosen blinked at 0.5 Hz for the duration of the TO. A new trial was then initiated. Choosing the holes associated to P1 and P2 throughout the session resulted in more pellets earned at the end of the session, compared to the P3 and P4 options. Thus, P1 and P2 were “optimal choices” while P3 and P4, leading to longer TO punishment and less pellets earned, were “non-optimal choices”. Risk-related decision-making was indexed with the choice score (% optimal choices - % non-optimal choices). Responses during the 5sec ITI were scored as premature responses, and signaled by a 5sec TO period before the initiation of a new trial. The percentage of premature responses [(#premature responses / #trial initiated) * 100] was an index of motor impulsivity. After 20 free-choice sessions of rGT, saline was injected i.p. 30min before sessions to control for the stress of injections, for a minimum of 5 sessions. Three consecutive saline sessions were selected to represent the baseline performance of the rats during rGT. A mixed factorial analysis of variance (ANOVA) was performed on the 3 selected days for baseline performance, with line (i.e. RHA or RLA) as between-line factor, and choice score over the 3 selected days as within-subject factor. The 3 days selected for baseline were considered stable if no main effect of session nor line x session interaction were observed. When stable baseline was observed, the effect of the activation or inhibition of mPFC-NAc pathway on motor impulsivity and risk-related decision-making was assessed in a single-session of rGT performed 30min after the administration of CNO (1mg/kg, ip).

### Tissue preparation and Immunohistochemistry

At the end of the experiments, rats received a lethal dose of sodium pentobarbital (150mg/kg; i.p.; Streuli Pharma AG, Uznach, Switzerland) and were transcardially perfused with saline 0.9% followed by 4% paraformaldehyde (PFA, Sigma-Aldrich, Steinhem, Germany) in phosphate buffer saline (PBS). Brains were extracted and post-fixed in PFA for 24 hours, before being placed in 30% sucrose in PBS for 48-72 hours for cryoprotection. Then, the brains were embedded in optimal cutting temperature compound (CellPath, Newtown, Powis, United Kingdom), frozen in isopentane, and stored at -80°C until processing. Brains were then cut into 40 μm slices along the caudo-rostral extension of the mPFC and NAc. Free-floating slices were stained with Hoescht (Abcam, #ab828550), and mounted with Fluorsave reagent (Merck KGaA, Damstadt, Germany). Fluorescent images were acquired using a widefield fluorescence slide scanner microscope (Zeiss Axioscan Z1, Gottingen, Germany) and analyzed using Zen 2.0 software (Zeiss, Gottingen, Germany).

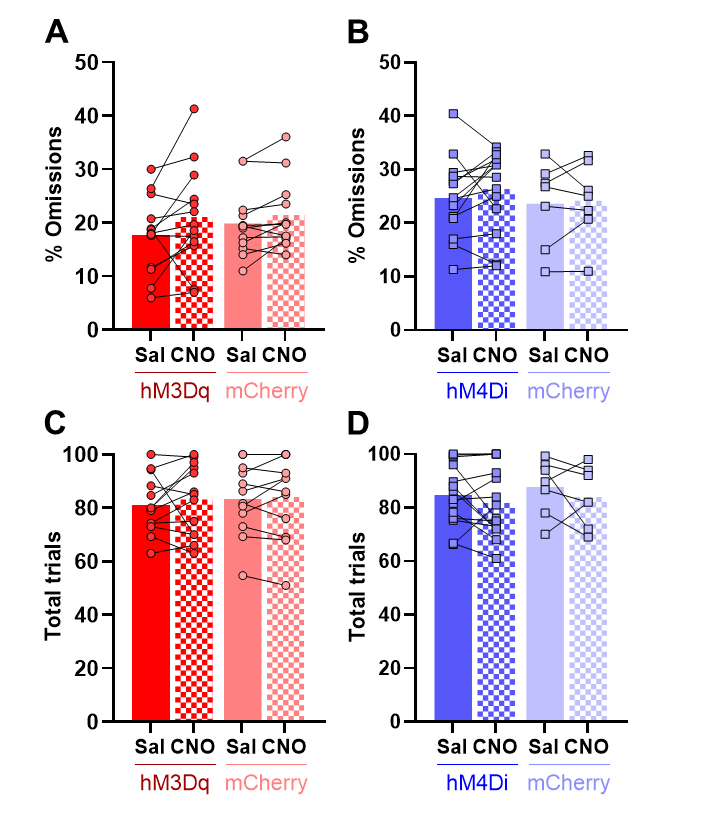

**Supplementary Fig. S1**: Effect of the modulation of mPFC-NAc pathway on rGT performances. **(A)** Compared to saline, CNO treatment increased omissions in RHA-hM3Dq and in RHA-mCherry groups. **(B)** In RLA rats, CNO had no effect in RLA-hM4Di or in RLA-mCherry groups. Relative to saline, CNO treatment had no effect on the total number of trials between **(C)** RHA-hM3Dq or RHA-mCherry groups and between **(D)** RLA-hM4Di or RLA-control groups.
